# Palmitoylation of PD-L1 Regulates Its Membrane Orientation and Immune Evasion

**DOI:** 10.1101/2021.10.17.464736

**Authors:** Siya Zhang, Hongyin Wang, Zhongwen Chen, Ilya Levental, Xubo Lin

**Affiliations:** Institute of Single Cell Engineering, Beijing Advanced Innovation Center for Biomedical Engineering, Beihang University, Beijing 100191, China; Institute of Basic Medical Sciences, Chinese Academy of Medical Sciences, Beijing 100005, China; Department of Molecular Physiology and Biological Physics, University of Virginia, Charlottesville, VA 22903, USA; Multiscale Research Institute of Complex Systems, Fudan University, Shanghai 200433, China; Key Laboratory of Ministry of Education for Biomechanics and Mechanobiology, School of Biological Science and Medical Engineering, Beihang University, Beijing 100191, China

**Keywords:** PD-L1 Palmitoylation, Raft Affinity, Membrane Orientation, Cancer Immunotherapy, Simulations and Experiments

## Abstract

Recently identified palmitoylation of PD-L1 is essential for immune regulation. To elucidate the underlying molecular mechanism, we performed giant plasma membrane vesicle (GPMV) experiments, µs-scale all-atom molecular dynamics (MD) simulations and immune killing experiments. GPMV experiments indicated that PD-L1 palmitoylation enhanced its lipid raft affinity. MD simulations revealed dramatically different membrane orientation states of PD-L1 in liquid-ordered (*L*_*o*_, lipid raft) compared to liquid-disordered (*L*_*d*_, non-raft) membrane environments. *L*_*d*_ region promoted the “lie-down” orientation of PD-L1, which could inhibit its association with the PD-1 protein on immune cells and thus promote the immune killing of cancer cells. This hypothesis was supported by immune killing experiments using *γδ*T cells as effector cells and NCI-H1299 lung cancer cells as target cells. In short, our study demonstrates that the palmitoylation affects PD-L1’s membrane localization and then membrane orientation, which thus regulates its binding with T cell PD-1 and the immune regulation. These observations may guide therapeutic strategies by explicating the regulation of immune checkpoint proteins by post-translational modifications and membrane environments.

## Introduction

In the past decades, cancer immunotherapy experienced unprecedented development in both the fundamental and clinical researches, revolutionizing the future of cancer treatment^[1-2]^. Among these approaches, immune checkpoint proteins such as programmed cell death protein 1 (PD-1) and programmed death-ligand 1 (PD-L1) have attracted great attention^[3-4]^. When PD-L1 on cancer cell membranes attaches to PD-1 on T cell membranes, it can effectively inhibit killing of cancer cells by T cells^[5-7]^. Hence, blocking binding between PD-L1 and PD-1 has become an effective way to boost the cancer-killing ability of T cells^[8-9]^. Especially, high-resolution structures of PD-1 and PD-L1 complex^[10]^ have stimulated the rapid development of various drugs including antibodies^[11]^, peptides^[12]^, small molecules^[13]^ and nanomedicines^[14]^ to directly block their binding. Also, the transfection of the PD-1 gene into cancer cells with highly expressing PD-L1 can also recover the immune responses of T cells^[15]^. This is because on the surface of cancer cells the expressed PD-1 can directly bind to PD-L1, which makes the membrane orientation of PD-L1 not suitable to attach PD-1 on T cell membranes. In other words, the membrane orientation thermodynamics of cancer cell PD-L1 is essential for their binding to T cell PD-1 and thus the immune regulation.

As shown in **Fig. 1a**, the full-length PD-L1 consists of an extracellular portion (position 19-238), a single α-helical transmembrane domain (position 239-259), and the cytoplasmic domain (position 260-290). The extracellular domain can be further divided into two functional Ig-like domains: V-type (position 19-127) and C2-type (position 133-225). Recently, Yang *et al*.^[16]^ identified a single palmitoylation modification site in its cytoplasmic domain at Cys272 with bioinformatics tools^[17]^. Palmitoylation is the covalent attachment of palmitic acid to typically cysteine (S-palmitoylation) of membrane proteins, which usually locates at the cytosolic part of the protein close to the inner cell membrane. They further found that site-specific mutation of the palmitoylation site or inhibition of the relevant palmitoyltransferase (ZDHHC9) could disrupt PD-L1 palmitoylation and restore T-cell killing ability to two kinds of breast cancer cells. This well validated the role of the palmitoylation in regulating PD-L1 stability and thus the immune response of T cells. Concomitantly, Yao *et al*.^[18]^ concluded a similar role of PD-L1 palmitoylation in regulating T-cell immune responses against cancer cells, via reduced ubiquitination of PD-L1 and escape from the subsequent degradation, which is regulated by electrostatic interactions between the cytoplasmic domain and the cell membrane^[19]^. While the impact of PD-L1 palmitoylation in immune regulation is validated, the potential impact of PD-L1 palmitoylation on its binding to PD-1 is still not clear.

**Figure 1.**
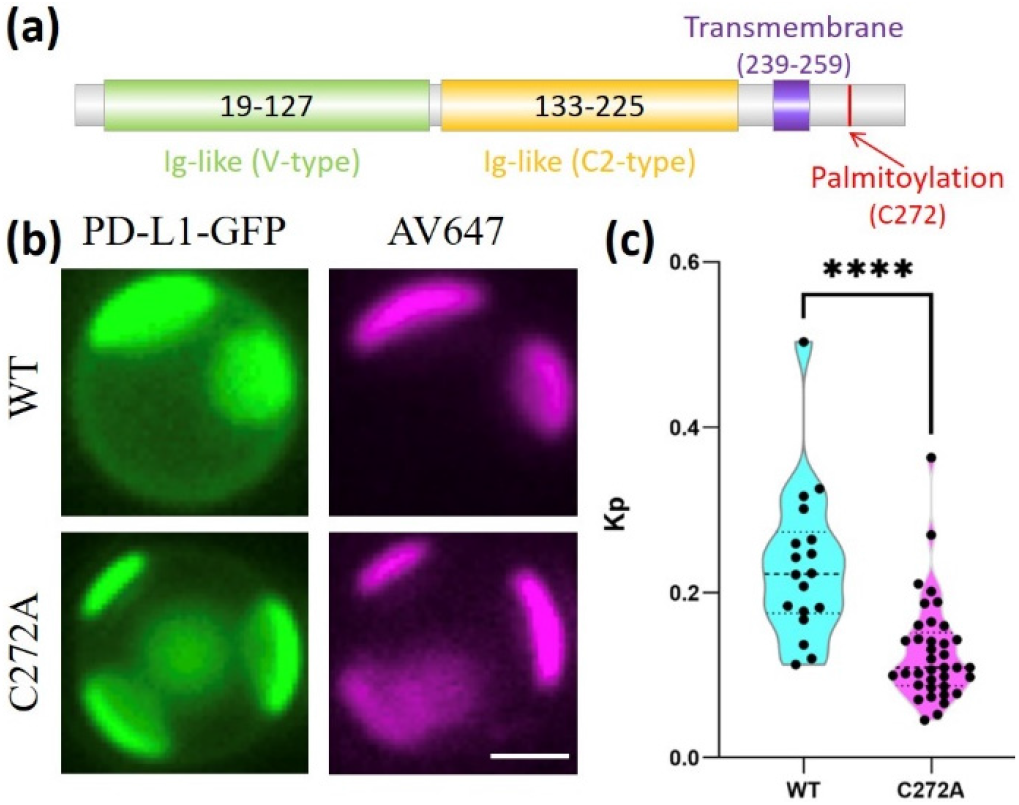
Palmitoylation promotes PD-L1’s lipid raft affinity. **(a)** Schematic of functional domains and palmitoylation site for full-length PD-L1 (290 aa). **(b)** The partitioning of wild-type (WT) and mutant (C272A) full-length PD-L1 in GPMVs. AV647 is a fluorescent probe for non-raft domain. Scale bar: 2μm. **(c)** *K*_*p*_ value for wild-type and mutant PD-L1.

One of the well-known roles for palmitoylation of transmembrane proteins is to enhance their affinity to lipid rafts^[20-23]^, which are nanoscale functional ordered membrane domains^[24]^. Because membrane orientation of transmembrane proteins can be modulated by the surrounding membrane lipid compositions^[25-27]^, we speculate that PD-L1 palmitoylation may promote its lipid raft affinity and thereby affect its membrane orientation, potentially affecting immune responses by modulating its interaction with PD-1. To test this hypothesis, we first performed giant plasma membrane vesicle (GPMV) experiments for wild-type PD-L1 (WT) and a palmitoylation-null mutant (C272A) to confirm the role of palmitoylation in determining PD-L1’s lipid raft affinity. Then, all-atom molecular dynamics (MD) simulations were used to reveal the relationship between membrane lipid composition and PD-L1’s membrane orientation. Last, we validated the implication from MD simulations via immune killing experiments using *γδT* cells and PD-L1 highly expressed NCI-H1299 lung cancer cells.

## Methods

### Giant Plasma Membrane Vesicle Experiments

GPMVs formation and *K*_*p*_ measurement were performed as previously described^[28-29]^. Briefly, MDA-MB-231 cells were transfected with PD-L1-GFP (Addgene #121139) or PD-L1-GPF-C272A. Then GPMVs were induced by incubating cells in GPMV buffer (150 mM NaCl, 10mM HEPES, 2mM CaCl2, 2 mM N-ethylmaleimide, pH 7.4) at 37°C for 3 h. To facilitate phase separation, 200 uM DCA (sodium deoxycholic acid) was added to the GPMV solution. Annexin V Alexa Fluor® 647 (AV647, ThermoFisher, #A23204; 1:100 dilution) was used to label non-raft regions on phase separated GPMVs. After cooling to <10 °C using Peltier stage (Warner Instruments), phase separated GPMVs were imaged using the epifluorescence microscope. The raft partitioning coefficient (*K*_*p*_) was calculated as 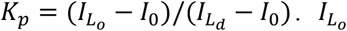 and 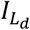 are the intensities in the protein channel from raft (AV647 poor) and non-raft (AV647 rich) regions. *I* is the background intensity from the protein channel.

### Molecular Dynamics Simulations

In our simulations, full-length PD-L1 without the signal peptide (position 19-290, **Fig. S1**) was used. Currently, the crystal structure for the transmembrane domain of PD-L1 is not available. The structure of its extracellular domain (PDB ID: 4Z18, 1.95 Å) is stable and conserved, while the cytoplasmic domain (PDB ID: 6L8R) is very flexible. Considering this, we employed the I-TASSER webserver^[30]^ using the crystal structure of PD-L1’s extracellular domain (PDB ID: 4Z18) as a template to obtain the full-length structure of PD-L1 (**Fig. S1**). In the current work, the liquid-ordered (*L*_*o*_) lipid membrane of PSM/POPC/Chol (0.61/0.08/0.31) and the liquid-disordered (*L*_*d*_) lipid membrane of PSM/POPC/Chol (0.23/0.69.0.08)^[31-32]^ were adopted to model the lipid raft and non-raft membrane domains in all all-atom MD simulations (PSM: palmitoylsphingomyelin, POPC: 1-palmitoyl-2-oleoyl-glycero-3-phosphocholine, Chol: cholesterol). CHARMM-GUI^[33-34]^ was employed to set up simulation systems of PD-L1 with or without palmitoylation embedded in *L*_*o*_ or *L*_*d*_ lipid membranes correspondingly. *L*_*o*_ membrane system consists of one palmitoylated PD-L1, 244 PSM, 32 POPC, 124 Chol, 47736 TIP3 waters, 150 mM NaCl, and its initial box dimension is 9.65nm×9.65nm×20.02nm. *L*_*d*_ membrane system consists of one PD-L1 without palmitoylation, 92 PSM, 276 POPC, 32 Chol, 58754 TIP3 waters, 150 mM NaCl, and its initial box dimension is 10.71nm×10.71×19.78nm.

CHARMM36m all-atom force field^[35]^ with grid-based energy correction maps for the protein conformational refinement^[36]^ and GROMACS 2019.6^[37]^ were used for all our MD simulations. We chose “cutoff” scheme for the Lennard-Jones potential, which was smoothly shifted to zero between 1.0 and 1.2 nm to reduce cutoff noise. Particle mesh Ewald electrostatics with a real-space cutoff of 1.2 nm was used. PD-L1, lipids and water&ions were coupled separately to Nose-Hoover heat baths^[38-39]^ with the coupling constant of 1 ps for T = 310 K. Pressure was kept at 1 bar using a semiisotropic Parrinello-Rahman coupling scheme^[40]^ with a coupling constant of 5 ps and a compressibility of 4.5×10^−5^ bar^-1^. LINCS algorithm was used to constrain bonds with hydrogen atoms. The nonbonded interaction neighbor list was updated every 20 steps with a cutoff of 1.2 nm. The simulations were run for 2 µs with the time step of 2 fs. System snapshots were rendered by VMD^[41]^.

In order to quantify the membrane orientation thermodynamics of PD-L1, we obtained the two-dimensional (2D) Gibbs free energy map with two reaction coordinates based the MD trajectories using the equation Δ*G*= −*RTln*(Ω/Ω_0_), where *R* is the gas constant (8.31 J⋅K^-1^⋅mol^-1^) and Ω is the density of states. One reaction coordinate (*ξ*_1_) is the center-of-mass distance along the membrane normal (z axis) between the distal sub-domain of PD-L1 (position 19-127, **Fig. 1a**) and the lipid membrane, while the other reaction coordinate (*ξ*_2_) represents the center-of-mass z-distance between the adjacent sub-domain of PD-L1 (position 133-225, **Fig. 1a**) and the lipid membrane. These two reaction coordinates can be directly calculated using GROMACS tool *gmx distance*^[37]^ over the MD trajectories.

### Human Peripheral Blood Mononuclear Cells Isolation and *γδ*T Cell Expansion

In order to obtain high-purity *γδ* T cells, we first collected peripheral blood samples from healthy volunteers with informed consent, which were approved by the ethical board of the Institute of Basic Medical Sciences at the Chinese Academy of Medical Sciences. Then, peripheral blood mononuclear cells (PBMCs) were isolated by Ficoll-Hypaque (Pharmacia, Sweden) density gradient centrifugation and cultured in RPMI-1640 medium (Gibco, USA) supplemented with 10% FBS and 200 IU/ml recombinant human interlukin-2 (IL-2) (Beijing Four Rings Biopharmaceutical, China), 5μM zoledronate acid (CTTQ Pharma, China). After 9 days of culture, *γδ* T cells of a purity >80% was used for the subsequent immune killing experiments.

### Cancer Cell Lines and Culture

NCI-H1299 human non-small cell lung cancer cells with high expression of PD-L1 were purchased from the American Type Culture Collection (ATCC) and cultured in RPMI-1640 medium (Gibco, USA) supplemented with 10% fetal bovine serum (FBS) at 37 °C in humidified 5% CO_2_. Trypsin (0.25%) EDTA (0.02%) solution (BI, Israel) was used for cell passage. Methyl-β-cyclodextrin (MBCD) was purchased from sigma Aldrich (USA) and used to extract cholesterols from NCI-H1299 cells according to the state-of-art protocol^[42]^.

### Flow Cytometry and Antibodies

In order to make sure the sufficient interactions of PD-1 and PD-L1 for our *γδ*T cells and NCI-H1299 cancer cells, flow cytometry was performed to detect the proportions of the expressions of PD-1 as well as *γδ*TCR for *γδ*T cells and of PD-L1 for NCI-H1299 cells. For cell surface staining, the *γδ*T cells and NCI-H1299 cancer cells were incubated with antibodies FITC anti-human gamma delta TCR (Clone B1.1, eBioscience ™, USA), PE/Dazzle ™ 594 anti-human PD-1 (CD279, Clone EH12.2H7, Biolegend, CA, USA), mouse anti-human PD-L1 (primary antibody, CD273, Clone OTI9E12, ORIGENE, MD, USA) and APC goat anti-mouse IgG (secondary antibody, Clone Poly4053, Biolegend, CA, USA) respectively for 30 min at 4 °C. Then, the cells were analyzed with ACEA NovoCyte™ (ACEA Biosciences, CA, USA), and the data were analyzed by NovoExpress software (ACEA Biosciences, CA, USA).

### *In Vitro* Cytotoxic Assay

NCI-H1299 cells as target cells were added to 96-well plates at a density of 5×10^5^ per well. The expanded *γδ*T cells as effector cells were incubated with target cells at an effector-to-target ratio of 5:1 and 10:1 overnight, and each condition was plated in quadruplicate. The three controls were background group, spontaneous release group and maximal release group. We detected the cytotoxicity by a lactate dehydrogenase (LDH) assay using the Cytotox 96 non-radioactive cytotoxicity assay kit (Promega, USA) according to the kit instructions. Cytotoxicity was calculated using the following formula:

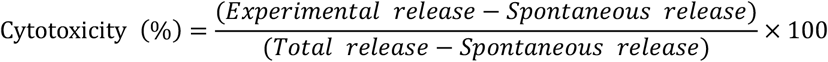

For each *γδ*T cells sample, the cytotoxicity of *γδ*T cells to NCI-H1299 cells treated with 0 mM, 1.25 mM or 2.5 mM MBCD are tested at least five times for each effector-to-target ratio. Considering individual differences in immunity, the relative killing efficiency was calculated by normalizing the cytotoxicity value with the average cytotoxicity value of 0 mM MBCD.

## Results and Discussion

### Palmitoylation Promotes PD-L1’s Lipid Raft Affinity

Multiple cancer cell lines including MDA-MB-231 were used in several independent experiments to validate the role of PD-L1 palmitoylation in the immune regulation ^[16, 18]^. Hence, in this work, we use MDA-MB-231 cells for our GPMV experiments to test whether PD-L1 palmitoylation affects its lipid raft affinity. For this purpose, we transfected wild-type (WT) and mutant (C272A) PD-L1 with a GFP tag into MDA-MB-231 cells and quantified their raft affinity in GPMVs. As shown in **Fig. 1b-c**, PD-L1 (green) is enriched in the liquid disordered membrane domain (marked by AV647), though with clear signal in the raft phase. Compared to WT PD-L1, the C272A mutant is nearly entirely excluded from the raft domain. Quantification of *K*_*p*_ indicated that palmitoylation indeed significantly enhanced the lipid raft affinity of PD-L1, which is consistent with our previous results for other transmembrane proteins (e.g. LAT - Linker for activation of T-cells)^[20-21, 43]^. Since the protein-lipid interactions play key roles in determining the dynamics and functions of proteins, the significant differences in lipid compositions for lipid raft and non-raft domains may induce dramatic different membrane dynamics of PD-L1, which could be important for its functions. However, it was worth mention that PD-L1 both with and without palmitoylation mainly resides in the lipid non-raft domain, which is different from LAT^[21, 43]^. This is probably due to the position differences of the palmitoylation site relative to the transmembrane domain of the protein. For the case of PD-L1, Cys272 is relative far from the transmembrane domain (**Fig. 1a**), which thus have the limited effects in determining the final lipid raft affinity of the protein.

### Lipid Raft Affinity Determines PD-L1 Membrane Orientation Thermodynamics

GPMV experiments revealed that PD-L1 palmitoylation significantly enhanced its lipid raft affinity, with depalmitoylated PD-L1 being completely excluded from raft domains. To investigate the possible structural consequences of this difference, we performed µs-scale all-atom MD simulations of PD-L1 in two membrane systems. One system is *L*_*o*_ lipid membrane embedded with a palmitoylated PD-L1, the other system is *L*_*d*_ lipid membrane embedded with a PD-L1 without palmitoylation. Three-component lipid membranes were used to simulate *L*_*o*_ (PSM/POPC/Chol = 0.61/0.08/0.31) and *L*_*d*_ (PSM/POPC/Chol = 0.23/0.69.0.08) lipid membranes^[31-32]^, which provides insights for the lipid raft and non-raft membrane domains with complex lipid compositions.

Considering the importance of protein’s membrane orientations in the binding between PD-1 (immune cells) and PD-L1 (cancer cells)^[10]^, we mainly focused on the differences of PD-L1’s membrane orientation thermodynamics in *L*_*o*_ and *L*_*d*_ lipid membranes. For this purpose, the center-of-mass distances along z axis between two sub-domains (position 19-127, position 133-225, **Fig. 1a**) of PD-L1 and the lipid membrane were chosen as the two reaction coordinates (*ξ*_1_, *ξ*_2_). States of these two reaction coordinates were recorded in real time over the whole 2 µs all-atom MD simulations (**Fig. S2**). Using the equation Δ*G*= −*RTln* (Ω/Ω_0_) and the highest density of states as Ω_0_, we could obtain the corresponding 2D free energy maps (**Fig. 2a**). The system configurations for typical states in **Fig. 2a** were shown in **Fig. 2b**. We could clearly find that the probabilities for the “stand-up” (“S”) and “lie-down” (“L”) states of PD-L1 are dramatically different in *L*_*o*_ lipid membrane from those in *L*_*d*_ lipid membrane. Generally, when palmitoylated PD-L1 (PD-L1-Pal) reside in *L*_*o*_ lipid membrane, it can have more “stand-up” states, which likely facilitate the engagement between cancer cell PD-L1 and T cell PD-1. For *L*_*d*_ lipid membrane, de-palmitoylated PD-L1 (PD-L1-noPal) possesses more “lie-down” states, which are not likely to allow PD-1/PD-L1 binding. Besides, the transition between “stand-up” and “lie-down” states is relatively easier for palmitoylated PD-L1 in *L*_*o*_ lipid membrane. In order to highlight the role of lipid compositions, we also studied the membrane orientation thermodynamics of de-palmitoylated PD-L1 in *L*_*o*_ lipid membrane. As shown in **Fig. S3**, de-palmitoylated PD-L1 showed even much more preference to the “stand-up” states. This indicates that the surrounding lipid compositions play key roles in determining the preference to different membrane orientation conformations of PD-L1, and the presence of the palmitoylation can promote the transition between “stand-up” and “lie-down” states (**Figs. 2a** and **S3**).

**Figure 2.**
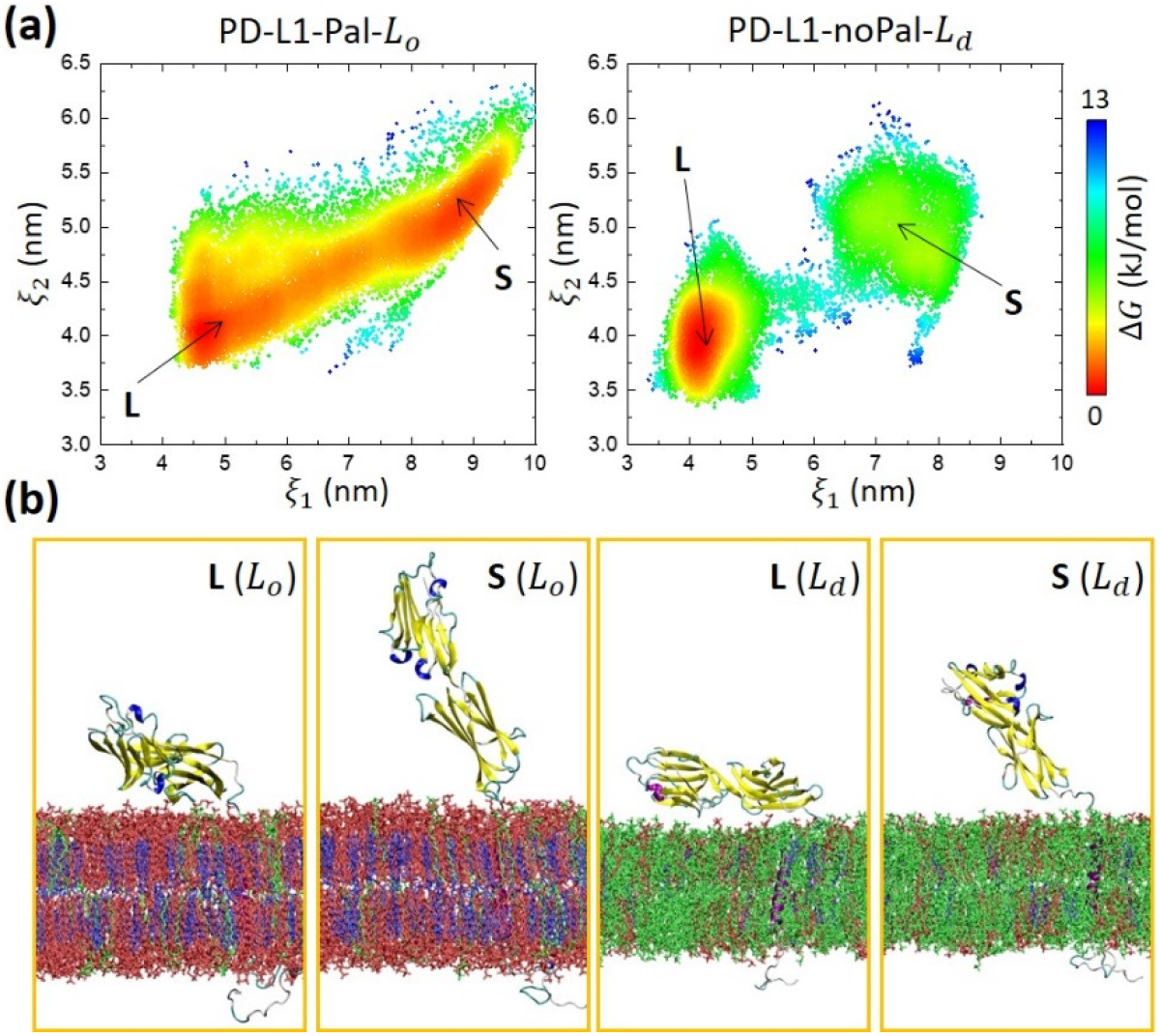
Membrane composition determines the orientation thermodynamics of PD-L1’s extracellular domain. **(a)** 2D free energy maps for the membrane orientation thermodynamics of PD-L1’s extracellular domain in *L*_*o*_ and *L*_*d*_ lipid membranes. “S” and “L” represents “stand-up” and “lie-down” states respectively. **(b)** Representative conformations indicated in **Fig. 2a**. PD-L1 is shown as carton and colored according its secondary structure, while its palmitoylation is visualized in *vdW* beads. PSM, POPC and Chol are colored as light red, green and blue bonds.

Different lipid compositions can induce different membrane orientation conformation of both membrane-bound^[26]^ and transmembrane proteins^[27]^. However, the mechanisms underlying these effects are often unclear. To determine whether direct interactions between lipids and the extra-membrane domain of PD-L1 was essential to the conformational distributions we observed, we simulated only the extracellular part of PD-L1 (position 19-235), which was anchored to the membrane via an ectopic C-terminal lipid anchor (palmitoylation at P235C). We then placed this truncated PD-L1 close to *L*_*o*_ or *L*_*d*_ lipid membranes, with the lipid anchor inserted into the membrane. The two new membrane systems were simulated for 1 µs and the corresponding 2D membrane orientation free energy maps are shown in (**Fig. 3**). While interactions between the extracellular domain of PD-L1 and *L*_*o*_/*L*_*d*_ membrane lipids have some effects on the transitions between various states, the PD-L1 extra-membrane domain behaved significantly different from the full-length PD-L1 (**Fig. 2**). Truncated PD-L1 could experience both “stand-up” and “lie-down” states easily both in *L*_*o*_ and *L*_*d*_ lipid membranes. Thus, considering the flexibility of the cytoplasmic domain of PD-L1, we propose that the significant effects of lipid environment on PD-L1’s membrane orientation thermodynamics (**Figs. 2** and **3**) probably originate from the transmembrane domain of PD-L1.

**Figure 3.**
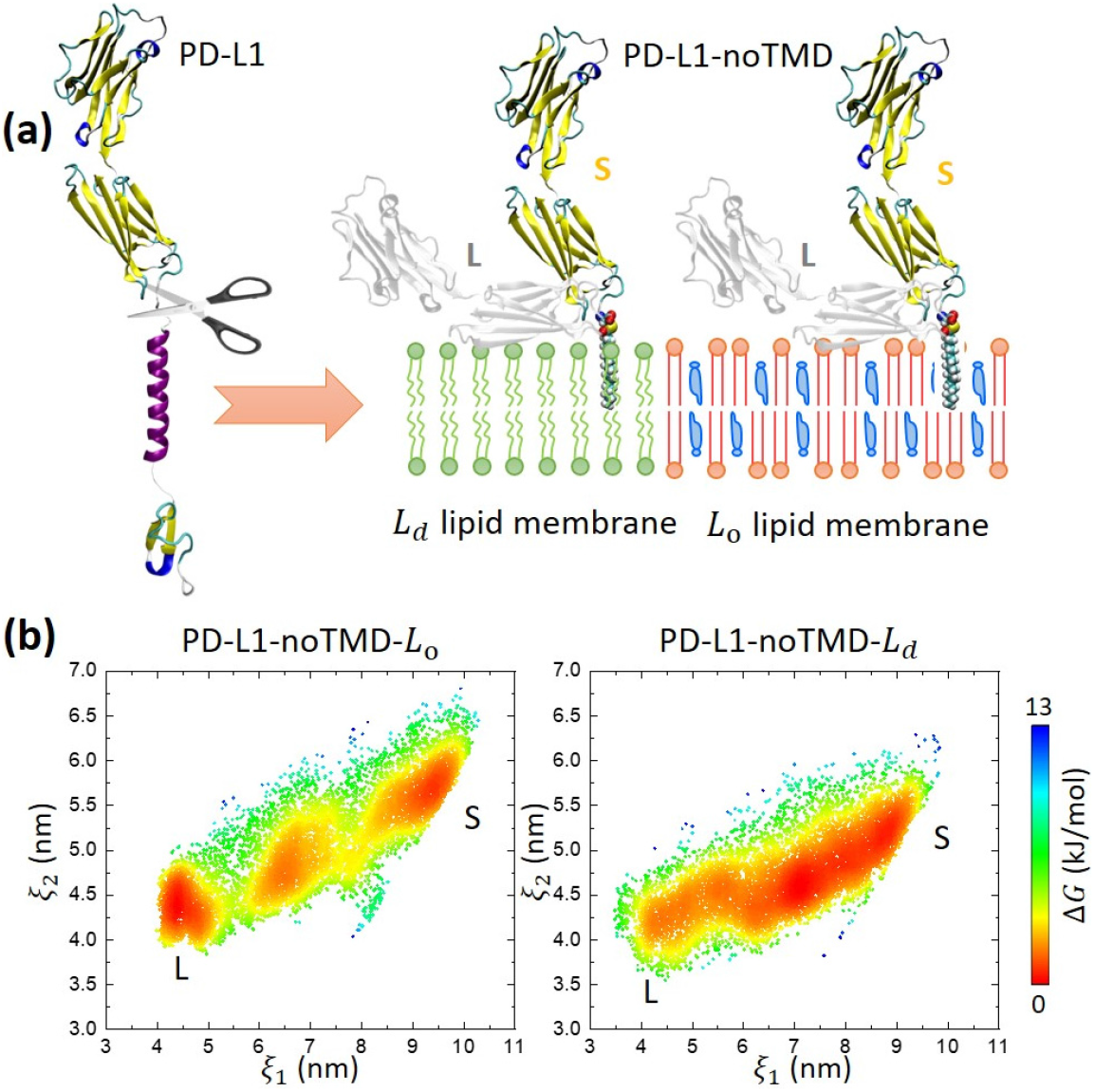
The absence of PD-L1’s transmembrane and cytoplasmic domains induces dramatically different orientation thermodynamics of its extracellular domain. **(a)** The transmembrane and cytoplasmic domains of PD-L1 are truncated and replaced with a lipid anchor (palmitoylation). The truncated PD-L1 is then placed in *L*_*o*_ and *L*_*d*_ lipid membranes for quantify its orientation thermodynamics transitioning between “stand-up” (S, multicolor) and “lie-down” (L, gray). **(b)** 2D free energy maps for the membrane orientation thermodynamics of truncated PD-L1 in *L*_*o*_ and *L*_*d*_ lipid membranes.

### Cholesterol Depletion from Cancer Cells Promotes *γδ*T Cell Killing Efficiency

PD-L1 palmitoylation promotes its lipid raft affinity and changes its membrane orientation thermodynamics. PD-L1 in lipid raft domain can access the “stand-up” state easier than in non-raft domain. Thus, we hypothesized that palmitoylation/raft association would promote the binding of PD-1 and PD-L1 and thereby suppress cytotoxicity of immune cells against cancer cells. Since cholesterol is required for lipid rafts, we tested the effects of lipid rafts on PD-L1 function by depleting membrane cholesterol from cancer cells via methyl-beta cyclodextrin (MBCD). We chose NCI-H1299 lung cancer cells and *γδ*T cells to perform immune killing experiments (**Fig. 4a**). *γδ*T cells have the advantage of tissue tropism and early activation, especially the intrinsic immune-like MHC non-restrictive recognition, allowing them to directly recognize a broad spectrum of tumor-associated antigens and exert cytotoxic activity. NCI-H1299 cells have a relatively high expression of PD-L1^[44]^ (**Fig. 4b**), while *γδ*T cells will have sufficient expressions of PD-1 and *γδ*TCR under the stimulation of cancer cells^[45-47]^ (**Fig. 4c**). MBCD was used to extract cholesterol molecules from NCI-H1299 cancer cell membranes. It is suggested that the MBCD concentration not higher than 3 mM is suitable for deleting cholesterols without significantly reducing the viability of cells^[42]^. Hence, we tested the situations of 0 mM, 1.25 mM and 2.5 mM MBCD in our immune killing experiments. These three cases are supposed to have increased cholesterol deleting ability. For the quantification of the killing efficiency of *γδ*T cells to NCI-H1299 cells, LDH assay was used. As shown in **Fig. 4d** and **Fig. S5**, the killing efficiency of *γδ*T cells to NCI-H1299 cells treated with 2.5 mM MBCD is significantly enhanced for all the tested samples. In other words, deleting the cholesterols of NCI-H1299 cells can recover the immune killing ability of *γδ*T cells, which is consistent with and can be elucidated by the results of our MD simulations. The cell membrane will change from cholesterol-rich (*L*_*o*_-like) to cholesterol-poor (*L*_*d*_-like) by deleting the cholesterols. PD-L1 in the *L*_*d*_-like cell membrane prefers the “lie-down” state, which inhibits the binding of PD-1 with PD-L1 and thus promote the immune normalization.

**Figure 4.**
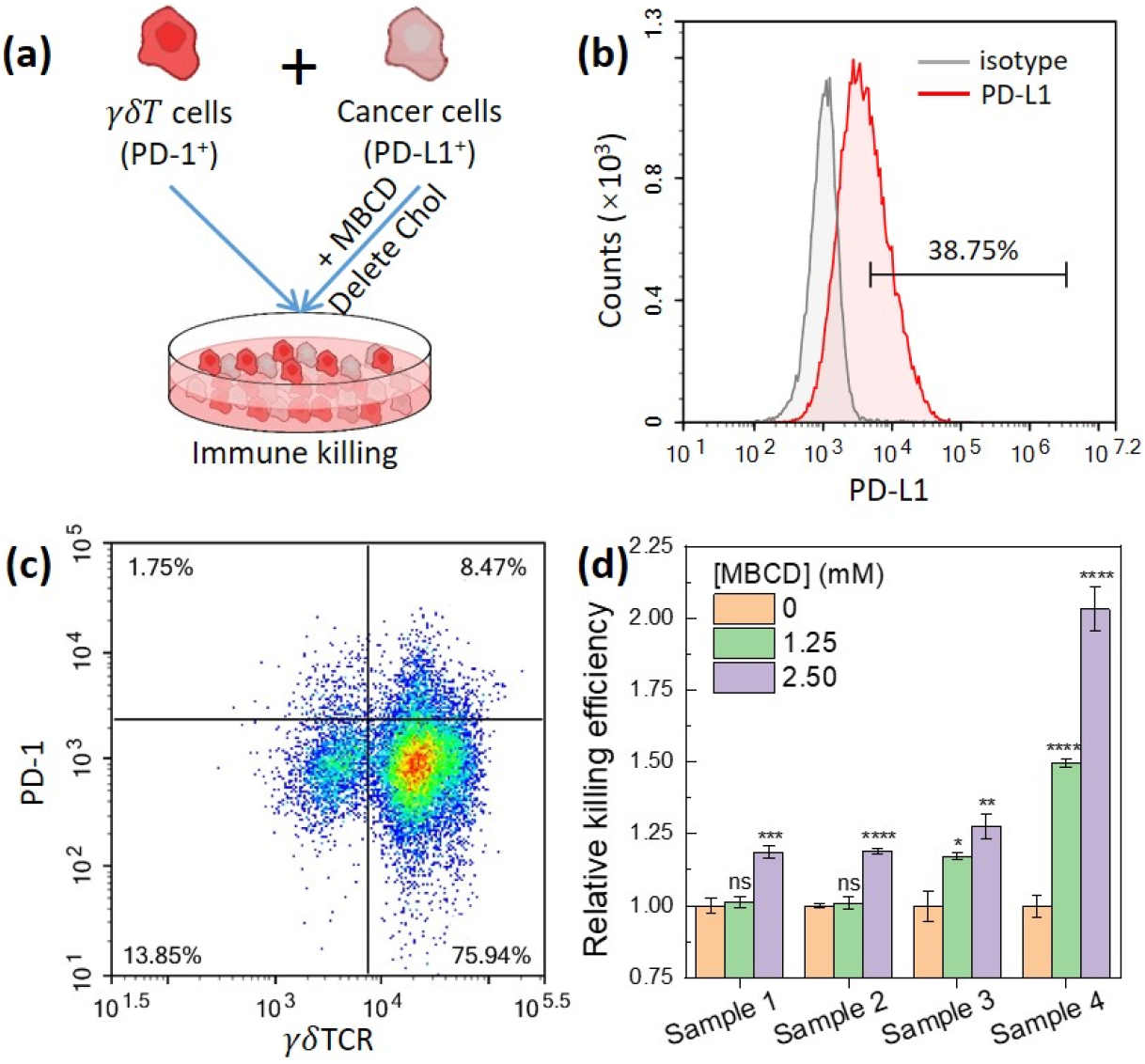
Removal of cholesterol molecules from cancer cells enhances the killing efficiency of immune cells. **(a)** Schematics of immune killing experiments based on immune checkpoint proteins PD-1 and PD-L1. **(b)** Flow cytometry analyzing the expression of PD-L1 in NCI-H1299 lung cancer cells. Number above the bracketed line indicates percent PD-L1^+^ cells. **(c)** Flow cytometry analyzing the expression of PD-1 and *γδ*TCR in *γδ*T cells. Numbers in quadrants indicate percent cells in each throughout. **(d)** Relative killing efficiency of *γδ*T cells over NCI-H1299 lung cancer cells with the treatment of MBCD of different concentrations with the effector-to-target ratio of 5:1.

## Conclusions

In this work, we proposed a new physical mechanism for the roles of PD-L1 palmitoylation in immune regulation by a combination of cellular experiments and atomistic simulations (**Scheme 1**). GPMV experiments confirmed that PD-L1 palmitoylation enhanced the protein’s lipid raft affinity. µs-scale all-atom MD simulations then revealed the membrane orientation thermodynamics of PD-L1 extracellular domain in *L*_*o*_ and *L*_*d*_ lipid membranes, indicating that PD-L1 in *L*_*d*_ lipid membranes prefers the “lie-down” state, which we hypothesized would inhibit the binding of PD-1 with PD-L1, thus inhibiting immune suppression. This was validated in immune killing experiments of *γδ*T cells to NCI-H1299 lung cancer cells whose membrane cholesterol were depleted using MBCD. In short, roles of PD-L1 palmitoylation in the immune regulation can be summarized in two points (**Scheme 1**). On the one hand, PD-L1 palmitoylation can prevent the ubiquitination and its subsequent protein degradation process^[18]^. On the other hand, PD-L1 palmitoylation can enhance its lipid raft affinity (**Fig. 1**) and the probability of the “stand-up” membrane orientation state (**Figs. 2**), which thus promote the binding of PD-1 with PD-L1 and suppress the killing of immune cells to cancer cells (**Fig. 4**).

**Scheme 1.**
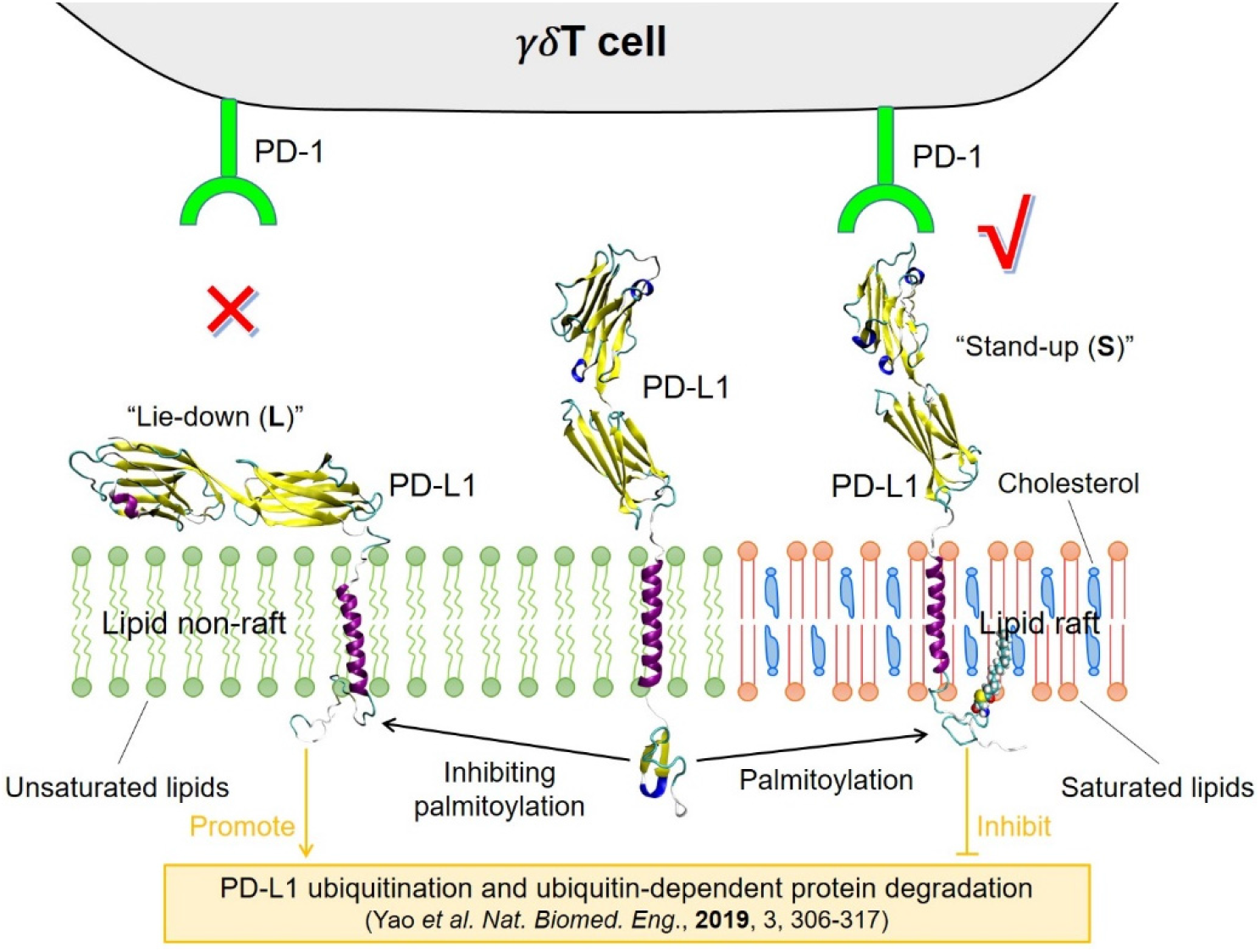
Two key roles of PD-L1 palmitoylation in the immune regulation: **(1)** PD-L1 palmitoylation inhibits its ubiquitination and the corresponding ubiquitin-dependent protein degradation (orange part)^[18]^. **(2)** PD-L1 palmitoylation enhances its lipid raft affinity and changes its membrane orientation thermodynamics, which promotes its binding with PD-1 and thus suppresses the immune responses of *γδ*T cells (This work).

## Supporting Information

Additional supplementary figures.

## Acknowledgments

This work was supported by the National Natural Science Foundation of China (*No*. 21903002, XL), the Fundamental Research Funds for the Central Universities (*No*. YWF-20-BJ-J-632, XL), the Open Fund of State Key Laboratory of Membrane Biology (*No*. 2020KF09, XL) and US National Institutes of Health/National Institute of General Medical Sciences (R35 GM134949, R01 GM124072, IL). We are grateful to Center for High Performance Computing of Beihang University (BHHPC) for generous computing resources.

## TOC Graphics

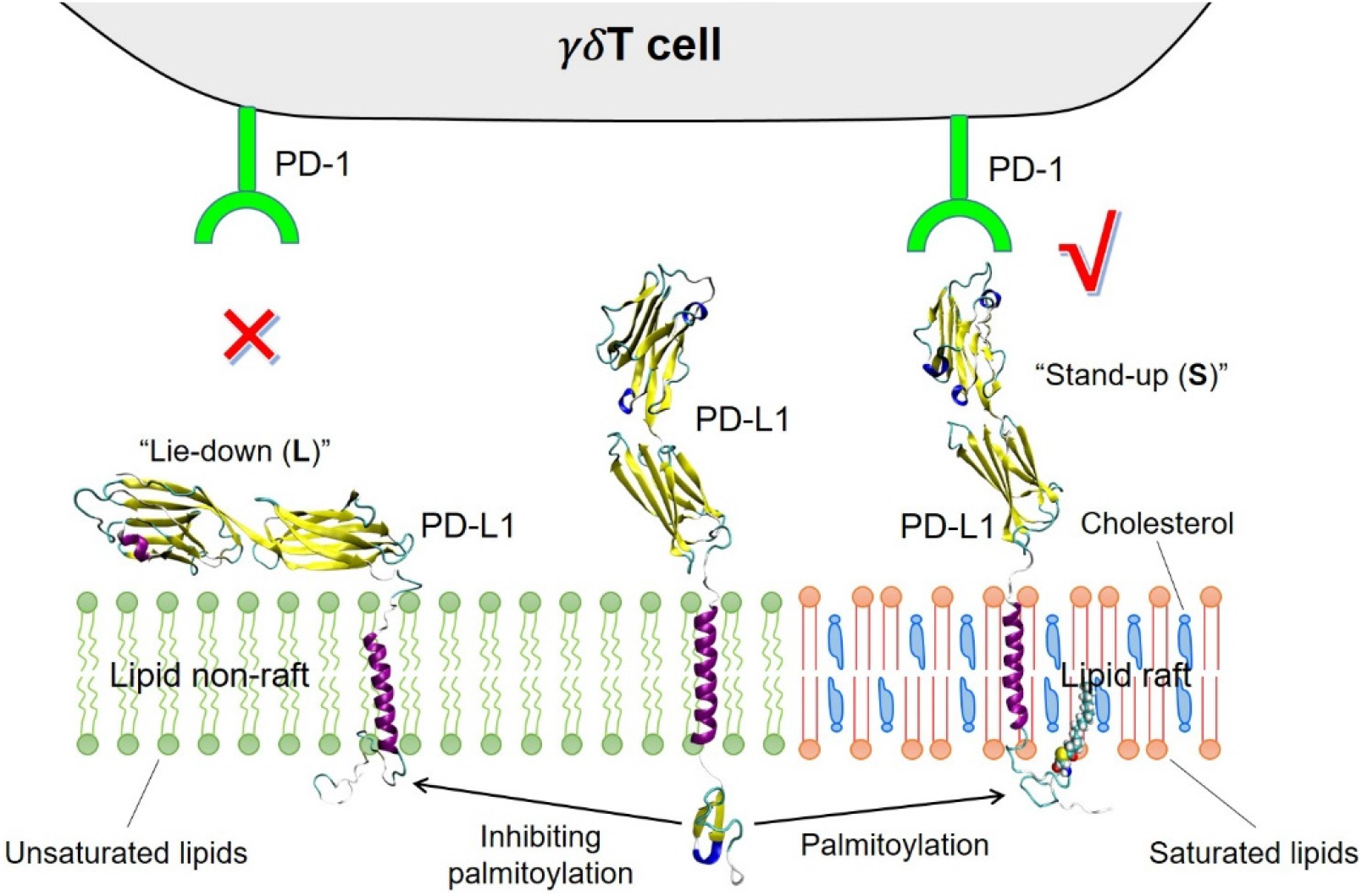

PD-L1 palmitoylation regulates immune responses of T cells via changing its membrane orientation.

## Supporting Information

**Figure S1.**
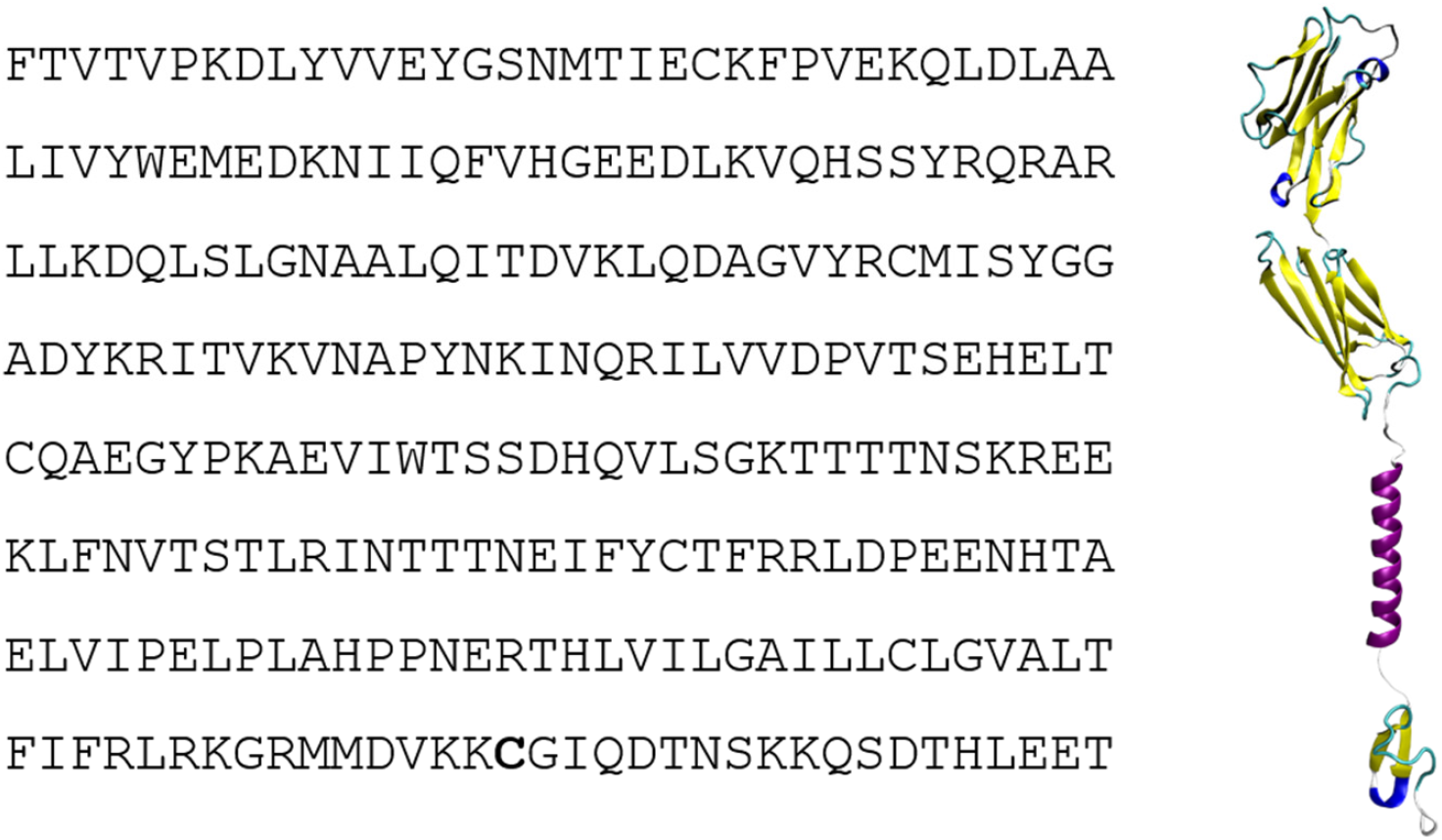
Amino acid sequence for full-length PD-L1 (left) and its corresponding molecular configuration (right). The palmitoylation site is shown in bold.

**Figure S2.**
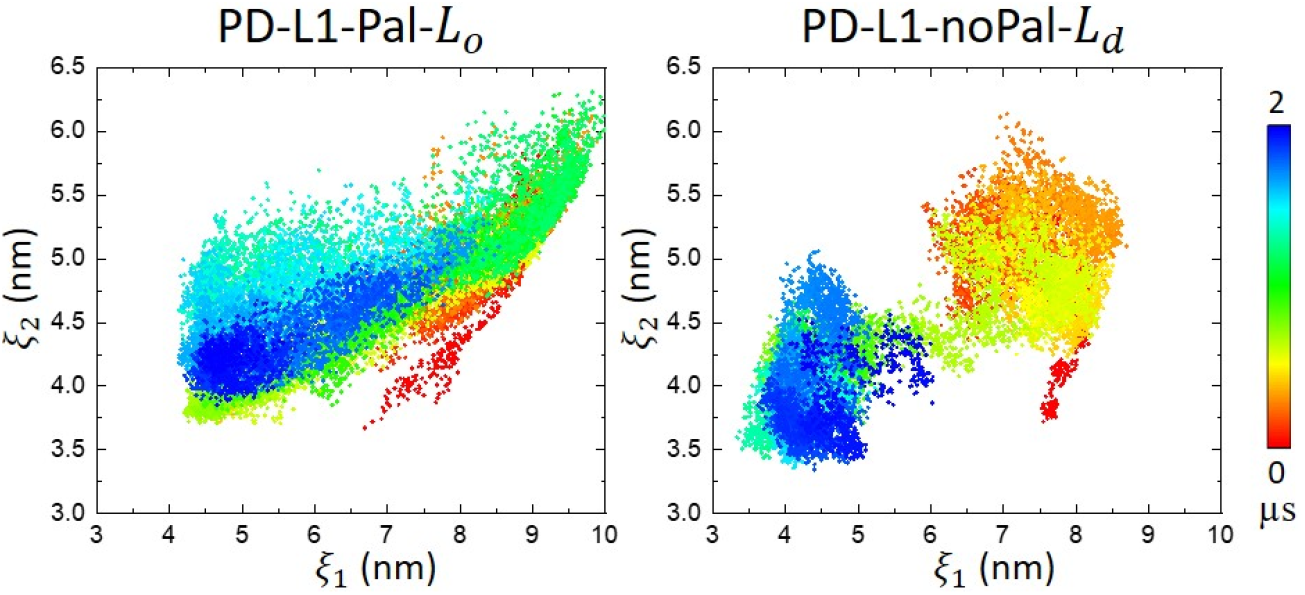
Time evolution of membrane orientation of full-length PD-L1 in *L*_*o*_ (PD-L1-Pal) and *L*_*d*_ (PD-L1-noPal) lipid membranes.

**Figure S3.**
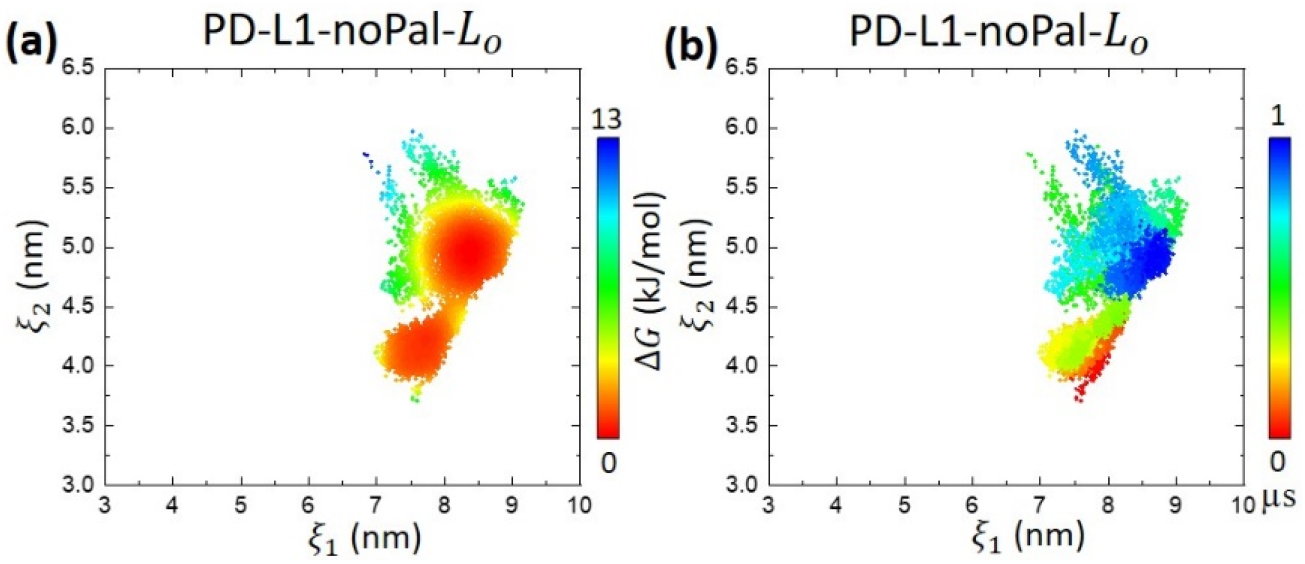
2D free energy map **(a)** and time evolution **(b)** of membrane orientation of PD-L1 without palmitoylation in *L*_*o*_ lipid membrane.

**Figure S4.**
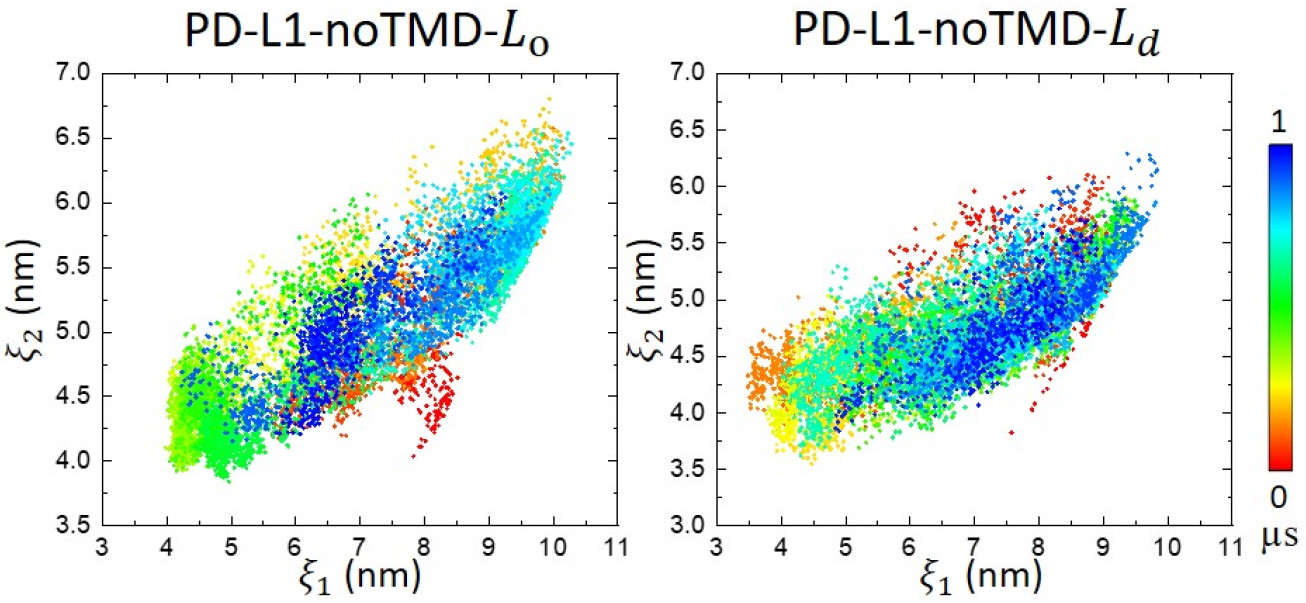
Time evolution of membrane orientation of PD-L1 without TMD in *L*_*o*_ and *L*_*d*_ lipid membranes.

**Figure S5.**
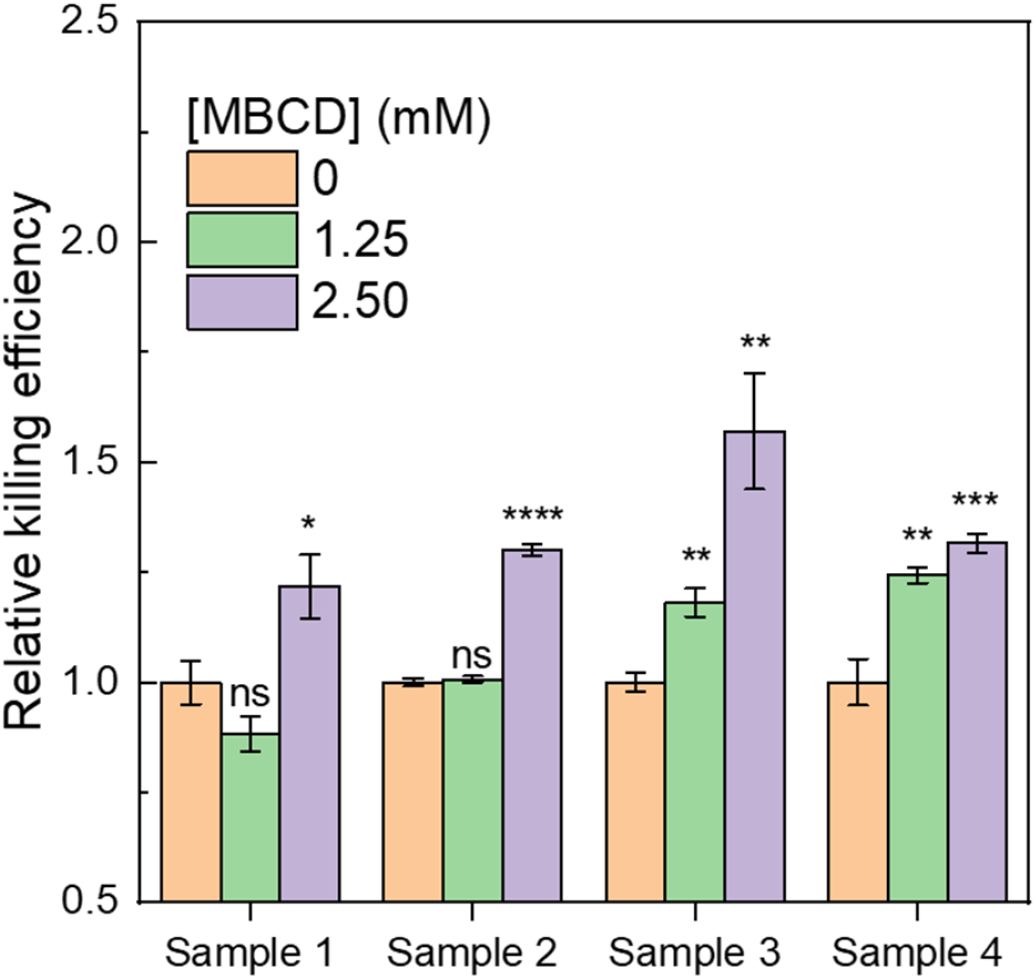
Relative killing efficiency of *γδ*T cells over NCI-H1299 lung cancer cells with the treatment of MBCD of different concentrations with the effector-to-target ratio of 10:1.

